# Elevated IL-1 beta plasma levels, altered platelet activation and cardiac remodeling lead to moderately decreased LV function in Alzheimer transgenic mice after myocardial ischemia and reperfusion

**DOI:** 10.1101/2025.09.04.674210

**Authors:** Lili Donner, Simone Gorressen, Jens Fischer, Margitta Elvers

## Abstract

**Introduction:** Neurodegeneration and dementia are key factors in Alzheimer’s disease (AD). The deposition of amyloid-ß into senile plaques in the brain parenchyma and in cerebral vessels known as cerebral amyloid angiopathy (CAA) are the main clinical parameters of AD. Acute myocardial infarction (AMI) and AD share a comparable pathophysiology. In detail, AMI survivors have a higher risk of vascular dementia and AD patients with AMI face a poor prognosis with a high rate of major adverse cardiovascular events. High blood levels of amyloid-ß40 are identified in patients with high risk for cardiovascular death. However, the underlying mechanisms and the consequences of AMI in AD patients are unclear to date.

**Methods:** AD transgenic APP23 mice were analysed in an experimental AMI using the closed-chest model.

**Results:** APP23 mice displayed significantly decreased left ventricular function as detected by FS/MPI (fractional shortening/myocardial performance index) after 24 h and 3 weeks after ligation of the LAD compared to WT controls. No differences have been observed in infarct and scar size. However, the analysis of cardiac remodeling after 3 weeks showed an altered composition of the collagen tissue of the scar with elevated tight but reduced fine collagen in APP23 mice. Altered scar formation was accompanied by elevated degranulation of platelets following activation of the collagen receptor GPVI.

**Conclusion:** The here presented results suggest that AD patients are at higher risk for cardiac damage after AMI that might increase the risk for cardiovascular death. This implies the need of a personalized therapy of AMI in AD patients.

## 1 Introduction

Alzheimer’s disease (AD) is a neurodegenerative disease and the most common form of dementia with a progressive decline in cognitive function (Clarfield, 2003). Beside age, major risk factors are a positive family history, hypertension and hypotension, high cholesterol levels, low levels of physical activity, obesity and the presence of epsilon 4 allele of the apolipoprotein E gene (APOE4) (Huang et al., 2004, Kivipelto et al., 2001, Kivipelto et al., 2005, Wallin et al., 2013).

AD patients present a specific neuropathological profile: The deposition of extracellular amyloid-ß (Aß) into senile plaques and the formation of intracellular neurofibrillary tangles (NFTs) that arise from hyperphosphorylated tau proteins (Long and Holtzman, 2019). More than 80% of AD patients develop cerebral amyloid angiopathy with Aß accumulation and aggregation in cerebral vessels (Selkoe, 2011, Thal et al., 2008). In addition, neuroinflammation and blood-brain-barrier dysfunction (Thal et al., 2008) induce neuronal loss and cognitive decline in AD patients.

AD and AMI share different pathophysiological hallmarks (Liu et al., 2024). AMI is associated with an increased risk of dementia induced by chronic hypo-perfusion of the brain after AMI caused by impaired LV function (Zuccalà et al., 2001). Furthermore, the subsequent release of emboli to the brain might be associated with dementia in AMI patients. Thus, AMI is associated with ischemic stroke with in turn increased the risk of dementia (Corraini et al., 2017, Sundbøll et al., 2016). Common risk factors such as diabetes mellitus, hypercholesterolemia, hypertension, inflammation and atherosclerosis suggest that AMI and AD may be independent but convergent diseases (Sundbøll et al., 2018, Thorp et al., 2022). Different studies suggest that there is a higher risk of dementia in AMI survivors (Sundbøll et al., 2018).

Platelets are the smallest blood cells and major regulators of hemostasis and thrombosis, but are also a key factor in the pathology of AMI (Krott et al., 2024, Reusswig et al., 2023, Reusswig et al., 2022) and AD (Chen et al., 1995, Donner et al., 2016, Donner et al., 2020, Gowert et al., 2014, Jarre et al., 2014, Koupenova et al., 2018, Leiter and Walker, 2020, Li et al., 1998, Wolska et al., 2023). Platelet dysfunction is associated with AMI (Michaels et al., 2000, Reusswig et al., 2022) and several neurodegenerative diseases such as AD (Beura et al., 2022, Jarre et al., 2014, Leiter and Walker, 2020). Hyper activation of platelets has been observed in experimental models of AMI (Reusswig et al., 2022) and AD (Jarre et al., 2014). In aged transgenic mice modeling Alzheimer’s disease (APP23) with parenchymal plaques and CAA, pre-activated platelets adhere to vascular amyloid-β deposits, leading to cerebral vessel occlusion (Gowert et al., 2014, Jarre et al., 2014). Thus, platelet activation might be another key element in both, AMI and AD.

In this study, we examined the consequences of ischemia and reperfusion injury in APP23 mice. We detected elevated IL1-β plasma levels and platelet degranulation after activation of the collagen receptor GPVI at 21 days after LAD ligation. Furthermore, moderate but significantly decreased LV function was associated with adverse cardiac remodeling while infarct and scar size were not affected.

## 2 Materials and Methods

### 2.1 Animals

The transgenic AD mouse model APP23 on a C57BL/6J background was used as described elsewhere (Donner et al., 2016, Sturchler-Pierrat et al., 1997). Food and water were provided ad libitum. APP23 and non-transgenic littermates (wildtype, WT) mice were fed standard chow. Mice of both sexes were used at an age of 22-25 month, where Aß pathology was pronounced including amyloid plaques in the parenchym and cerebral vessels. All experiments were performed in accordance with the German Law on the protection of animals and approved by a local ethics committee (LANUV, North-Rhine-Westphalia, Germany, reference number: reference numbers AZ 84-02.04.2013.A473, AZ 81-02.05.40.21.041 and O 86/12).

### 2.2 Flow cytometry (platelet activation and Annexin-V binding)

Heparinized whole blood from APP23 and WT littermates was used and diluted in Tyrode’s buffer (134 mM NaCl,12 mM NaHCO3, 2.9 mM KCl, 0.34 mM Na2HPO4, 20 mM HEPES, 10 mM MgCl2, 5 mM glucose, 0.2 mM CaCl2, pH 7.35) and washed twice. For the analysis of platelet activation, blood samples were mixed with fluorophore-labeled antibodies form Emfret Analytics (P-selectin exposure as marker for degranulation: Wug.E9-FITC, #M130-1; active form of α_IIb_β_3_ integrin: JON/A-PE, #M023-2) and 2 mM CaCl2 and stimulated with indicated agonists for 15 min at RT. For the analysis of pro-coagulant activity, Annexin-V binding to cells was determined by flow cytometry.

For determination of Aß binding to platelets, we used a FITC-labeled Aß antibody that detects APP and Aß at the surface of platelets.

### 2.3 Experimental model of acute myocardial infarction (AMI) and reperfusion in mice

A closed-chest model of reperfused myocardial infarction was used in order to reduce surgical trauma and subsequent inflammatory reaction from the intervention as described recently (Reusswig et al., 2022). Briefly, APP23 and WT littermates were anesthetized with with Ketamin (100 mg/kg body weight, Ketaset®, company: Zoetis, Malakoff, France) and Xylazin (10 mg/kg body weight, Xylazin, company: WDT, Ulft, The Netherlands) by a singular intraperitoneal (i.p.) injection before surgery. Euthanasia was performed by cervical dislocation. The left anterior descending artery (LAD) was ligated for 60 min. followed by reperfusion for 24h and 3 weeks as indicated. Coronary occlusion was achieved by gently pulling the applied suture tight until ST-elevation was detected by echocardiography (ECG). Thereafter, reperfusion was confirmed by resolution of ST-elevation. After 24 h of reperfusion, hearts were removed and stained with TTC/Evans Blue–solution to stain damaged cardiac tissue of the left ventricle (LV), separated in the area at risk (ischemic area) and the infarcted area. The ratios of the different areas were quantified digitally by video planimetry. To determine LV function after AMI, ECG was performed at different time points after I/R using Vevo 3100 ultrasound machine (VisualSonics Inc., Bothell, WA, USA) to measure different parameters, e.g., ejection fraction (%), cardiac output (mL/min), fractional shortening (FS, %), stroke volume (µL) and FS/MPI (myocardial performance index) (%) with corresponding software. In another set of experiments, reperfusion was performed for 3 weeks (21 days).

### 2.4 IL-1ß plasma levels

For quantification of IL-1 in the plasma of mice at 24h post ischemia and reperfusion, heparinized blood was centrifuged 10 min for 650 g to retrieve the plasma. The cytokine amount was measured by IL-1 enzyme-linked immunosorbent assay (ELISA; DuoSet Mouse IL-1ß/IL-1F2, R&D systems, Minneapolis, MN, USA) following the manufacturer’s protocol.

### 2.5 Collagen staining of cardiac tissue at 21 days after ischemia and reperfusion

First, scar size was determined by collagen staining 21 days after myocardial infarction by using Gomori’s one step trichrome staining. To this end, heart sections were prepared as mentioned before. The infarct size was expressed as the percentage of the total left ventricular (LV) area. Picrosirius red staining (Morphisto, Frankfurt am Main, Germany) was used for quantification of interstitial collagen deposition and Celestineblue-solution (Sigma, St. Louis, CA, USA) was used to stain nuclei. Interstitial collagen was measured in percent by area fraction.

To distinguish between tight and fine collagen, collagen density was analyzed by polarized light microscopy and evaluated by Image J software (Version number V 1.8.0, creator: Wayne Rasband, National Institutes of Health and the Laboratory for Optical and Computational Instrumentation LOCI, University of Wisconsin, (Madison, WI, USA)) as described recently (Reusswig 2022).

### 2.6 Statistical analysis

All statistical analyses were performed with GraphPad Prism (Prism 9; Graph Pad Software, Inc.). Unpaired t-test or one-way ANOVA, for two groups or three groups. The results were presented as the mean with each individual data point or in bar graph ± SEM. A p-value <0.05 was considered significant (*p < 0.05, **p < 0.01).

## 3 Results

### 3.1 LV function, infarct size, IL-1ß plasma levels and platelet activation in APP23 mice 24h after AMI

AD and AMI share common risk factors and might exacerbate disease pathology and all-cause mortality when patients suffer from both diseases. Therefore, we analyzed the consequences of AMI in the AD transgenic mouse line APP23. Experimental ischemia and reperfusion injury was induced in APP23 and WT control mice and LV function was investigated by echocardiography after 24 h of reperfusion. As shown in figure 1, cardiac output, fractional shortening, ejection fraction and stroke volume were not altered between APP23 and WT control mice (figure 1A-B, D-E). However, when we determined FS/MPI, we detected significantly reduced FS/MPI in APP23 mice compared to WT controls (figure 1C) suggesting reduced LV function in AD transgenic mice.

**Figure 1.**
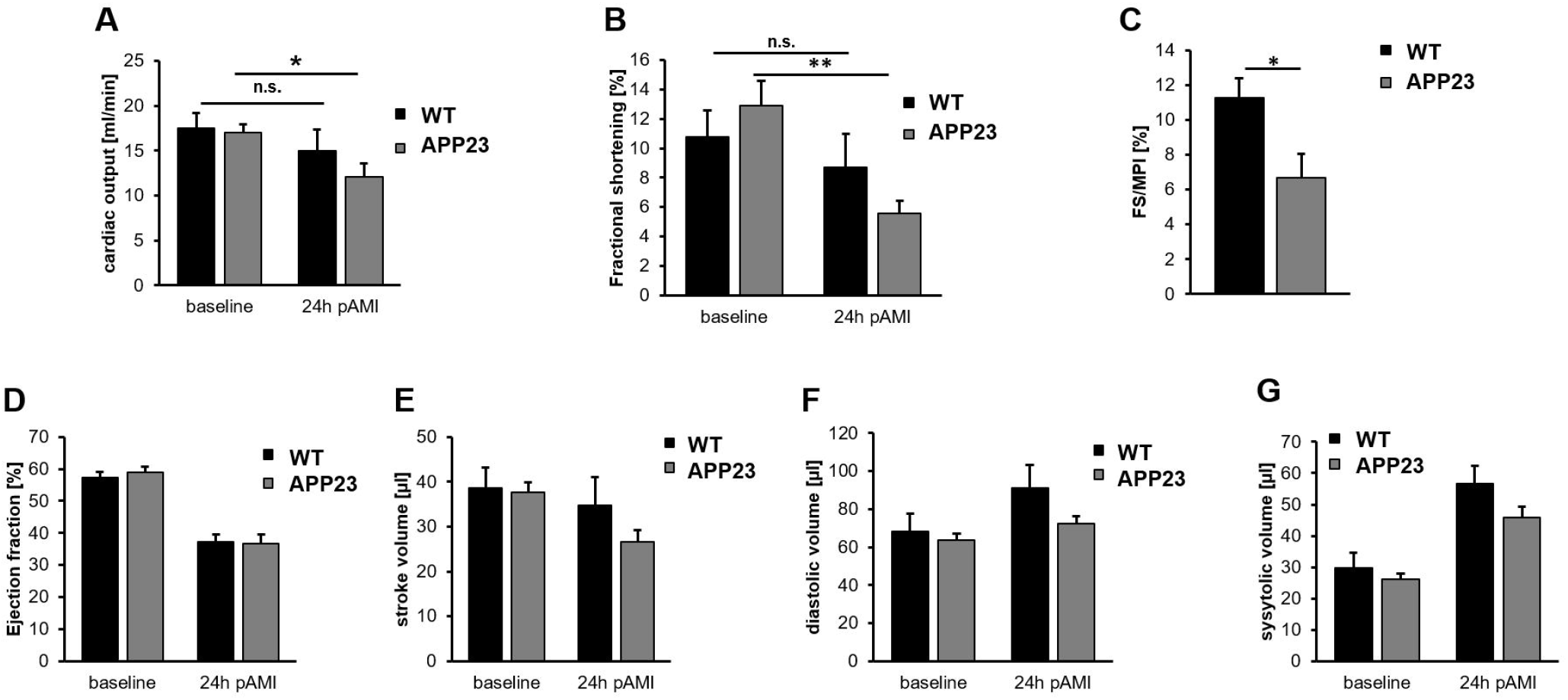
Moderately reduced left ventricular function in APP23 mice 24h post AMI. Echocardiographic analysis of cardiac function by determination of (A) cardiac output (ml/min.), (B) fractional shortening (%), (C) FS/MPI (%), (D) ejection fraction (%), (E) stroke volume (µl), (F) diastolic volume [µl) and (G) systolic volume (µl). Baseline vs. 24h after ischemia and reperfusion are shown. Bar graphs indicate mean values ± SEM. Statistical analyses were performed using a multiple unpaired t-test. N = 6. *p < 0.05; **p < 0.01. FS/MPI = fractional shortening/myocardial performance index; n.s. = not signficant.

In contrast, infarct size was not different between APP23 and WT controls (figure 2A). However, we detected elevated IL-1ß plasma levels in AD transgenic mice suggesting elevated inflammation in these mice (figure 2B). Flow cytometry was used to analyze platelet pro-coagulant activity by AnnexinV binding of platelets (figure 2C) and platelet activation by determination of active integrin αIIbβ3 and P-selectin exposure (marker for degranulation). As shown in figure 2 D-E, no differences were observed between APP23 and WT control mice (figure 2D-E).

**Figure 2.**
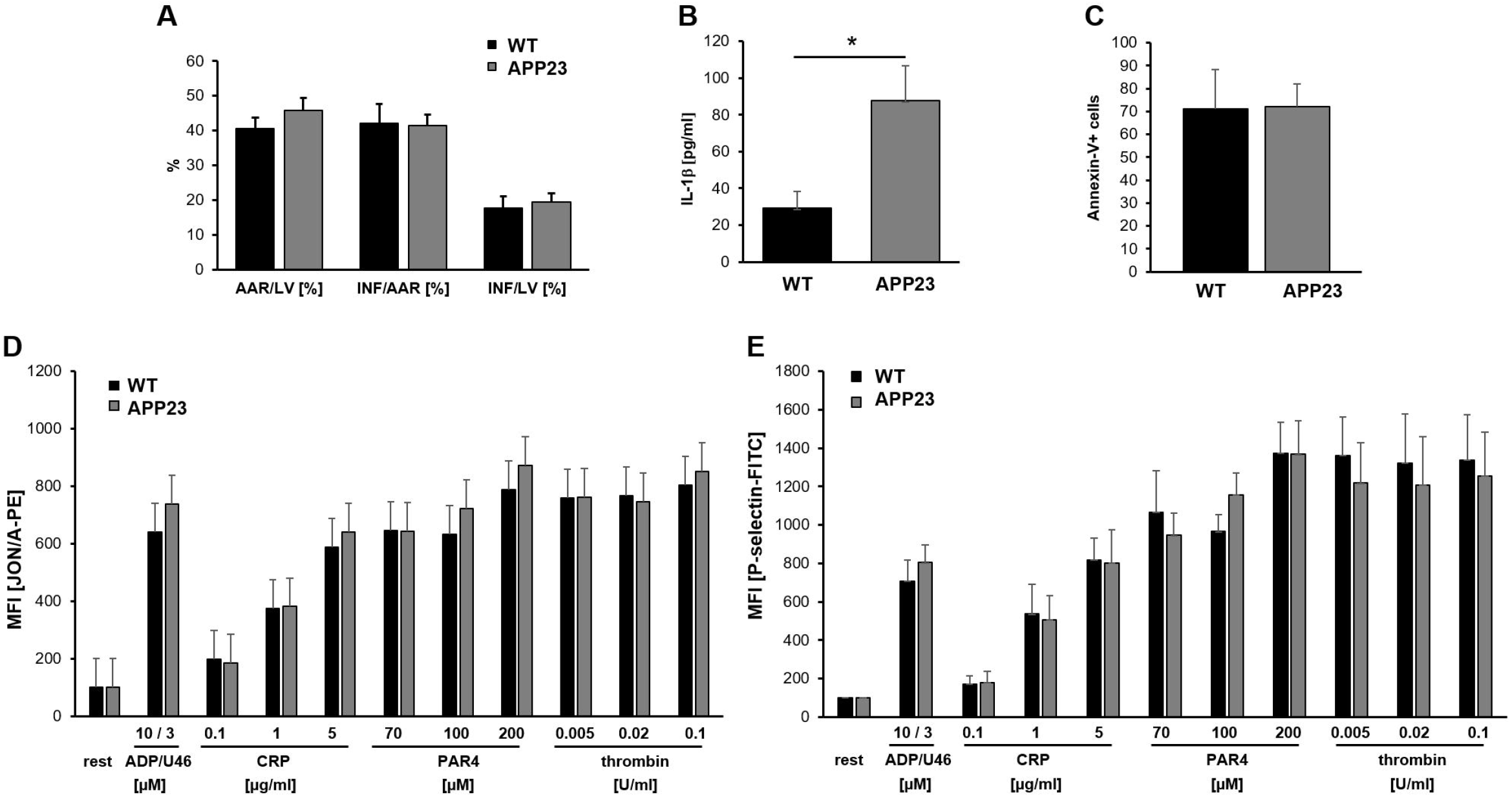
Unaltered infarct size and platelet activation but elevated IL-1β plasma level in APP23 mice 24h after ischemia and reperfusion. (A) Quantitative analysis of infarct size as the percentage of area at risk (% Inf/AAR, left panel) 24 h post AMI. (B) IL-1ß plasma levels in WT vs. APP23 mice 24h post ischemia and reperfusion. (C-E) Flow cytometry was used to determine AnnexinV binding to platelets as marker for pro-coagulant activity and platelet activation following stimulation of platelets with indicated agonists. Platelet activation was detected by measuring active integrin αIIbβ3 using the JON/A antibody and by determination of P-selectin exposure at the platelet membrane as marker for degranulation. Results are shown as MFI (mean fluorescence intensity). Bar graphs indicate mean values ± SEM. Statistical analyses were performed using an unpaired t-test. N = 6. ADP = adenosinediphosphate; U46 = thromboxane analogue; PAR4 = PAR4 peptide, activates the thrombin receptor PAR4; CRP = collagen-related peptide, activates the collagen receptor GPVI.

Moreover, blood cell counts such as the number of red and white blood cells and platelets were not altered, neither 24h nor 21 days post ischemia and reperfusion (Suppl.-figure 1).

### 3.2 Reduced LV function after 21 days following ischemia and reperfusion injury in APP23 mice

The analysis of LV function after 3 weeks of ischemia and reperfusion in APP23 mice revealed again reduced FS/MPI while no alterations were detected with regard to cardiac output, fractional shortening, ejection fraction and stroke volume as analyzed by echocardiography (figure 3). Moreover, diastolic volume and systolic volume were unaltered between APP23 and WT control mice. Thus, moderately decreased LV function in APP23 mice after 24 h and after 21 days after ischemia and reperfusion revealed prolonged cardiac dysfunction in AD transgenic mice after AMI.

**Figure 3.**
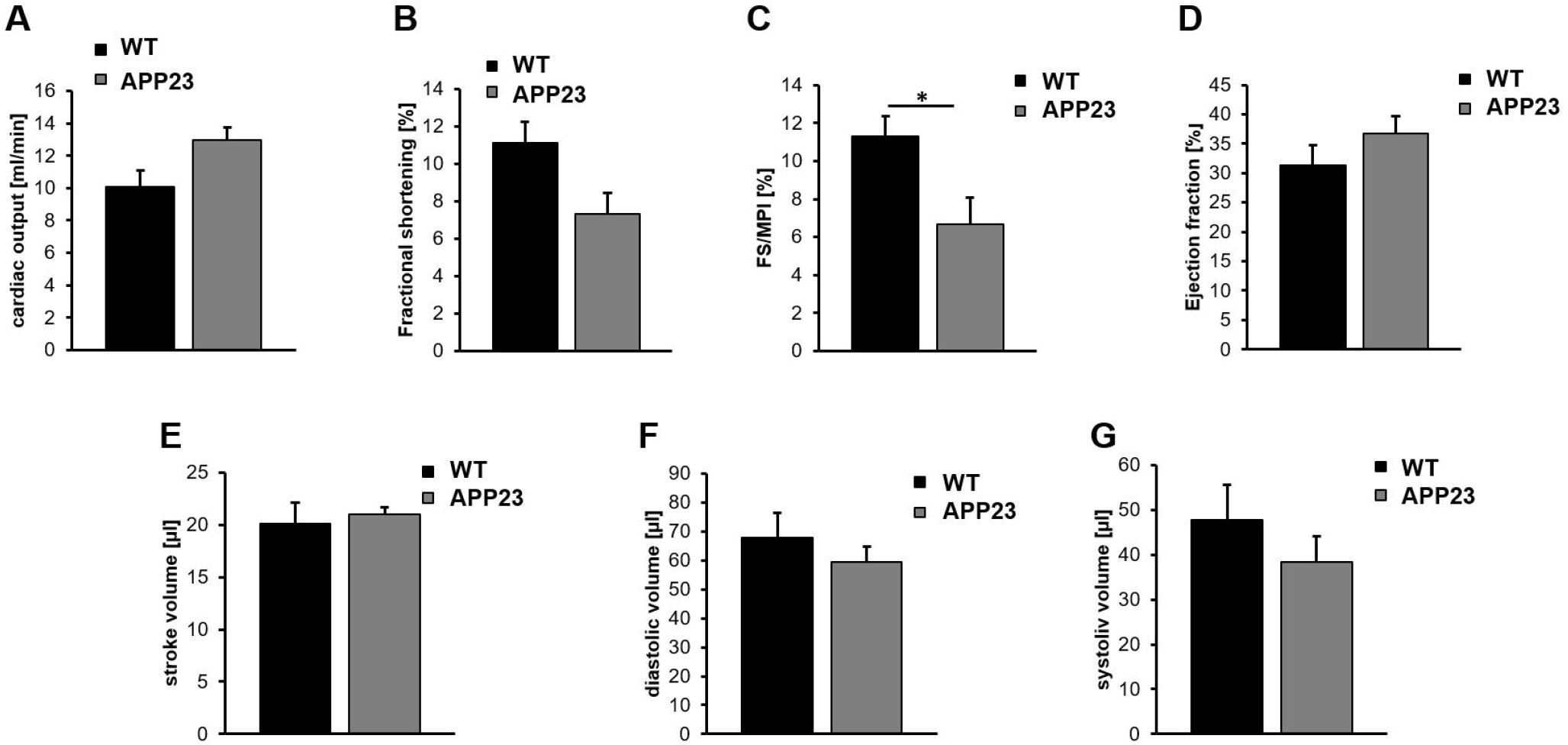
Moderately reduced FS/MPI in APP23 mice 21 days post ischemia and reperfusion. Echocardiographic analysis of cardiac function by determination of (A) cardiac output (ml/min.), (B) fractional shortening (%), (C) FS/MPI (%), (D) ejection fraction (%), (E) stroke volume (µl), (F) diastolic volume [µl) and (G) systolic volume (µl). WT vs. APP23 mice 21d after ischemia and reperfusion are shown. Bar graphs indicate mean values ± SEM. Statistical analyses were performed using a multiple unpaired t-test. N = 5-6. *p < 0.05. FS/MPI = fractional shortening/myocardial performance index.

### 3.3 Altered scar formation with elevated tight but reduced fine collagen in APP23 mice did not result in differences in scar size

21 days after ischemia and reperfusion injury, no differences in scar size were detected in AD transgenic mice (figure 4A-B). In addition, interstitial collagen was not altered (figure 4C). In contrast, cardiac remodeling as indicated by collagen composition of the scar revealed differences in the level of tight and fine collagen. In detail, increased levels of tight collagen but reduced fine collagen has been detected in APP23 compared to WT control mice (figure 4D-E). Thus, differences in the collagen composition might change scar quality but not scar size leading to reduced LV function in AD transgenic mice.

**Figure 4.**
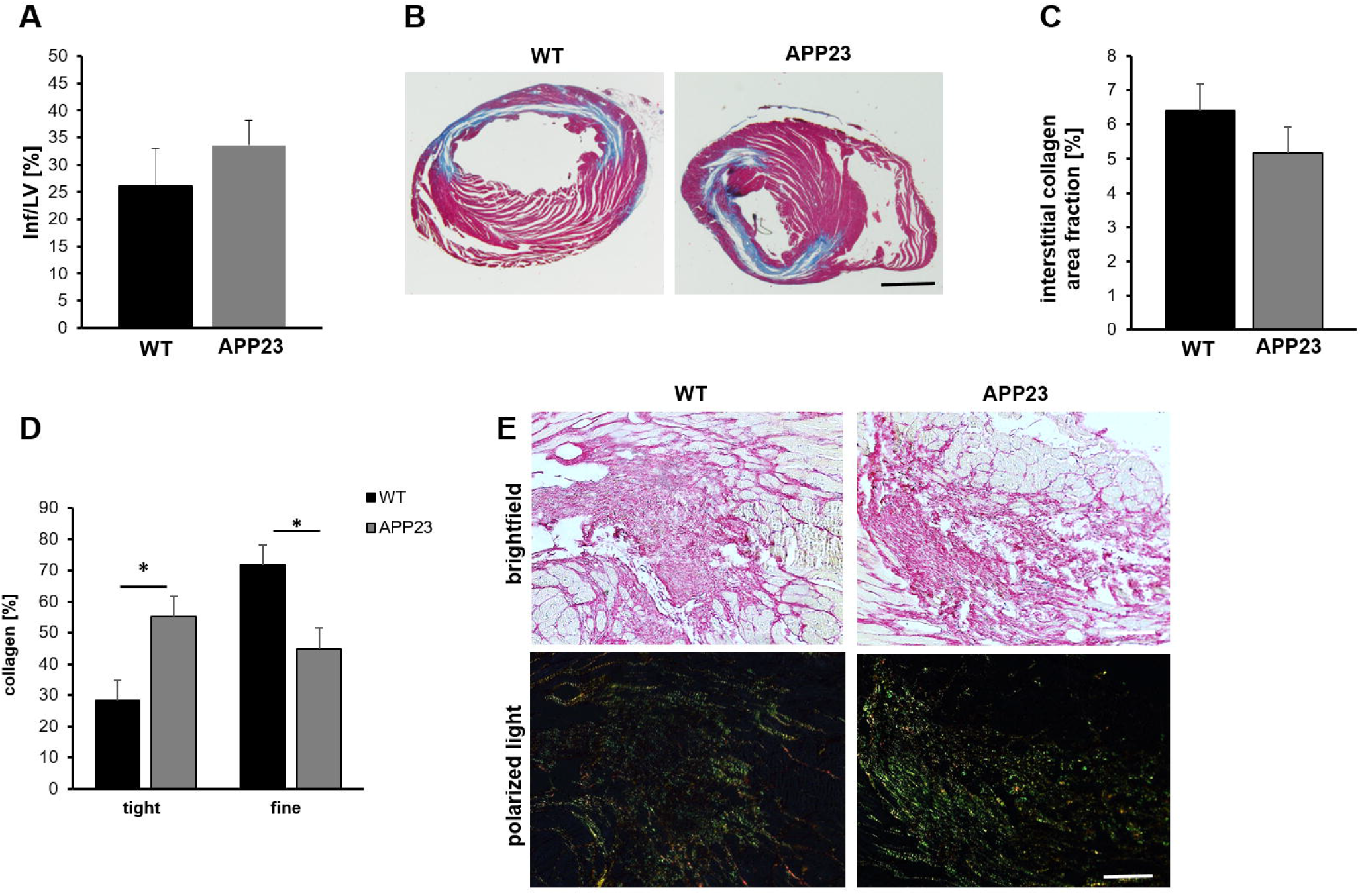
Unaltered scar size but different scar quality at 21 d post AMI. Quantification (A) and representative images of scar size (B) using Gomori’s trichrome staining 21 days post AMI. Infarcted area is stained in blue and healthy tissue is stained in red. Data are presented as means ± SEM. (C) Quantification of sirius-red staining of interstitial collagen in the remote zone of APP23 mice after 21 days of reperfusion compared to WT controls, (D) Analysis of collagen composition (D) and representative images of sirius-red staining (E) of APP23 mice after 21 days of reperfusion compared to their littermate controls. In bright field microscopy of the LV after AMI, the cytoplasm is stained in yellow and collagen is stained in red. Polarized light microscopy was used to identify thin collagen fibers (collagen type III) in green and dense collagen fibers (collagen type I) in yellow-red. Scale bar = 2 mm (B), 100 µm (E). Bar graph depicts mean values ± SEM. Statistical analyses were performed using an unpaired t-test. N = 6.

### 3.4 Alterations in platelet activation in APP23 mice after AMI suggest an impact of platelets in cardiac damage

Next, we analyzed platelet activation since platelets play a role in AMI and AD. As shown in figure 5, integrin αIIbβ3 activation as reflected by JON/A binding to platelets revealed no differences between groups (figure 5A). In contrast, P-selectin exposure at the platelet surface as marker for platelet degranulation was elevated in response to collagen-related peptide that activates the major collagen receptor GPVI at the platelet surface. Moreover, thrombin stimulation of platelets resulted in reduced P-selectin exposure of platelets from APP23 mice (figure 5B). Since platelets and in specific the collagen receptor GPVI are known to modulate inflammation and cardiac remodeling after AMI –at least in mice-(Reusswig et al., 2023, Reusswig et al., 2022), our results point to an impact of platelets in scar formation and cardiac function in AD transgenic mice as reflected by altered levels of tight and fine collagen in the scar and reduced FS/MPI as detected by echocardiography.

**Figure 5.**
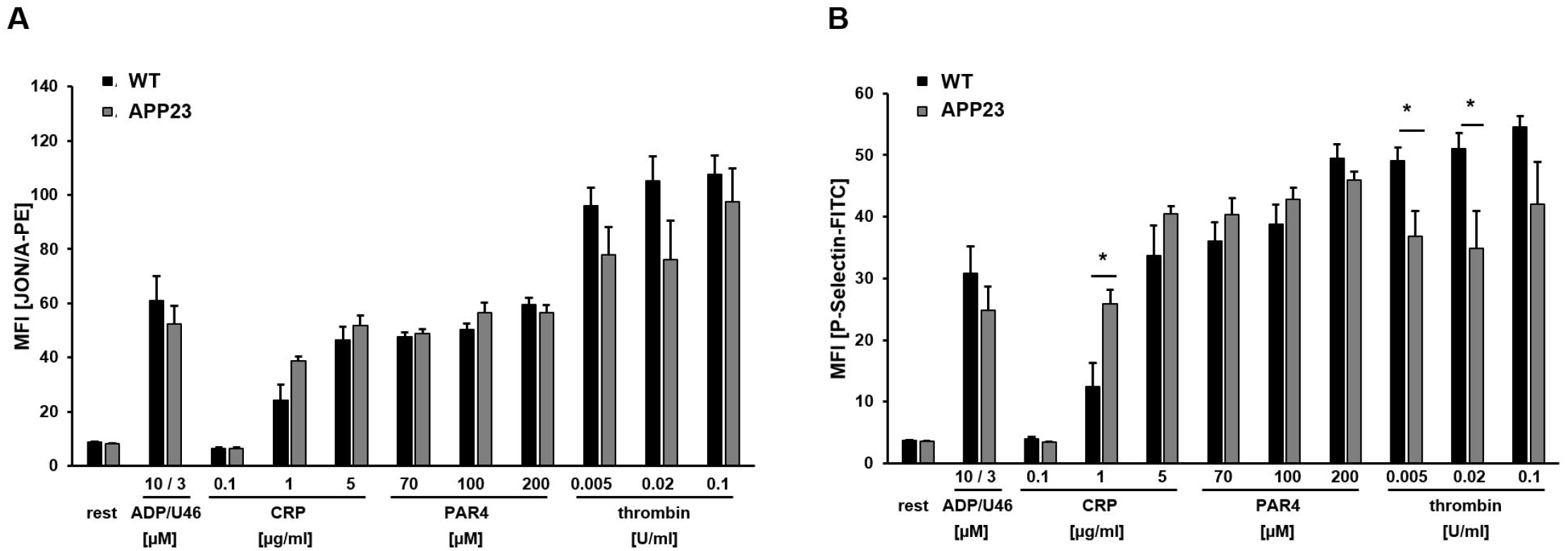
Flow cytometry analysis revealed enhanced platelet activation in response to GPVI in APP23 mice. (A) Active integrin (JON/A-PE binding to integrin αIIbβ3) and (B) platelet degranulation (P-selectin-PE) under resting and activated conditions were determined by flow cytometry using whole blood from APP23 and WT control mice. Different agonists were used to activate different signaling pathways in platelets. Data are represented as MFI. Bar graphs indicate mean values ± SEM. Statistical analyses were performed using a multiple unpaired t-test. *p < 0.05. ADP = adenosinediphosphate; U46 = thromboxane analogue; PAR4 = PAR4 peptide, activates the thrombin receptor PAR4; CRP = collagen-related peptide, activates the collagen receptor GPVI.; MFI, mean fluorescence intensity. N = 6.

## 4 Discussion

In recent years, an association of AD and heart failure including chronic and acute events such as AMI has been described. The multitude of published data focuses on the effects of cardiovascular disease on dementia including AD. Here, we provide evidence that mice with AD show impaired LV function and altered scar formation after ischemia and reperfusion injury. Thus, this is a first study to analyze the effects of AD including amyloid pathology in cerebral vessels and the brain parenchyma and how this affects cardiac damage after AMI, at least in mice. However, elevated IL-1β plasma levels and altered cardiac remodeling leads to only moderately decreased LV function as detected by reduced FS/MPI.

After AMI, echocardiography revealed only moderately reduced LV function in AP23 mice compared to controls at 24h and 21 days post ischemia and reperfusion injury. In detail, we detected significantly reduced FS/MPI whereas no differences were observed in cardiac output, fractional shortening, and ejection fraction and stroke volume. However, it has been shown that the FS/MPI ratio is the best noninvasive index of LV function in mice (Broberg et al., 2003). Moreover, MPI strongly correlates with dP/dt_max_ over a range of hemodynamic conditions in mice with the best correlation of FS/MPI with dP/dt_max_. Thus, our results clearly suggest that AD pathology is associated with a decreased LV function after AMI.

Published data describes a relationship between heart failure and AD. Thereby, heart failure as reflected by reduced cardiac output and neurohormonal activation, leads to reduced cerebral blood flow and hypoxia in the brain. Hypoxia results in tau hyperphosphorylation, amyloid precursor protein cleavage, dysfunction of the neurovascular unit and oxidative stress. These cellular responses then in turn induce the formation of neurofibrillary tangles and amyloid ß plaques, impaired amyloid-ß clearance and inflammation including dysfunction of microglia and thus AD (Cermakova et al., 2015). In line, Thorp and colleagues found an association of AMI and vascular dementia by inflammation induced by ischemia and reperfusion injury leading to neurovascular injury (Thorp et al., 2022). Also, the risk of vascular dementia is increased in survivors of AMI. The authors found that this association was even stronger in patients with stroke. However, Sundboll and colleagues were not able to provide evidence for an increased risk of AD in patients with AMI (Sundbøll et al., 2018).

Blood levels of Aß40 play a role in the survival rate of AMI patients. In detail, measuring the blood levels of Aß40 in patients with coronary heart disease may identify patients with high risk of cardiovascular death (Stamatelopoulos et al., 2015). Thus, Aß40 might serve as a biomarker to identify patients with cardiovascular disease and increased risk of death.

An interplay of cardiovascular risk factors and an increased risk of cognitive impairment and AD have been described by different groups. To this end, atherosclerosis plays a key role in this interplay including endothelial dysfunction and cerebral hypoperfusion. Moreover, diabetes mellitus (Craft et al., 1999), arterial hypertension (Guo et al., 2001) and dyslipidemia (Evans et al., 2004) are risk factors for cardiovascular disease and cognitive impairment (Tini et al., 2020). Different patient studies support this interplay of cardiovascular disease and AD. The prospective Rotterdam Study provides evidence for a two-three fold higher risk of developing AD in patients with severe atherosclerosis (Breteler et al., 1994). In a primary care patients study, this data was confirmed because coronary artery disease has an effect on mild cognitive impairment (MCI) in patients with AD (Bleckwenn et al., 2017).

Interestingly, carriers of the ApoE4 allele have shown to present a more marked association between coronary artery disease and neuropathological lesion of AD than patients who do not present this allele (Aronson et al., 1990. Thus, ApoE might affect both CAD and AD neuropathology. However, testing for ApoE cannot predict the development of AD in patients but might be useful to identify subjects at risk. Liu and colleagues found that AMI and AD share a comparable pathophysiology that maybe mediated by certain hub genes {Liu, 2024 #82). They identified the BCL6 gene to be essential for developing AMI and AD.

Recently, a retrospective cohort study provides evidence for patients with pre-existing AD to have a poor prognosis when they develop AMI because they showed a high rate of major adverse cardiovascular events (MACE) and all-cause mortality (Ozarek et al, EHJ 2024). Patients with AD were older, more likely to be female, and had a greater burden of individual comorbidities and multimorbidity. AD patients who were selected to undergo invasive management experienced reduce MACE and all-cause mortality compared to those managed conservatively. Thus, future research is important to identify patients who might benefit from invasive management. Furthermore, individualized multidisciplinary decision-making is key to provide the best therapy for AD patients with AMI.

Another link between AD and AMI might be platelets and their altered activation profile in both diseases. Platelets show a hyperactive profile in AD (Jarre et al., 2014) and affect CAA (Donner et al., 2016), at least in mice. In AMI, platelet activation is altered as well. In thrombocytopenic mice, a reduced inflammatory response was observed after ischemia and reperfusion injury (Reusswig et al., 2022). Moreover, platelets actively contribute to cardiac remodeling and scar formation and thus affect infarct size and heart function (Reusswig et al., 2023, Reusswig et al., 2022). Importantly, the major collagen receptor GPVI might play a dominant role in the above mentioned cellular responses. Thus, it is not surprising that increased platelet activation in APP23 mice was detected mainly following GPVI activation using platelets from mice at 21 days post ischemia and reperfusion. Therefore, we believe that platelets might be one important player in the relationship between AD and AMI.

Taken together, AMI in AD transgenic mice leads to elevated IL-1β plasma levels, increased platelet activation in response to GPVI and altered cardiac remodeling at 21 days post ischemia and reperfusion, resulting in moderately decreased LV function in APP23 mice. With regard to recently published data from AD patients with AMI, further research is important to further investigate the relationship of both diseases and to conduct a personalized therapy for patients at high risk of cardiovascular death.

## 6 Author contribution

ME and TS designed the study. LD and XX performed experiments. ME, TS and LD analyzed and interpreted data. ME wrote the manuscript with all authors providing feedback.

## 8 Acknowledgements

We thank Martina Spelleken and Dominik Semmler for providing outstanding technical assistance.

## 10 Data Availability Statement

The original contributions presented in the study are included in the article, further inquiries can be directed to the corresponding author.

