## Supplementary figures and images for "Elevated IL-1 beta plasma levels, altered platelet activation and cardiac remodeling lead to moderately decreased LV function in Alzheimer transgenic mice after myocardial ischemia and reperfusion"

### Supplemental Fig. 1

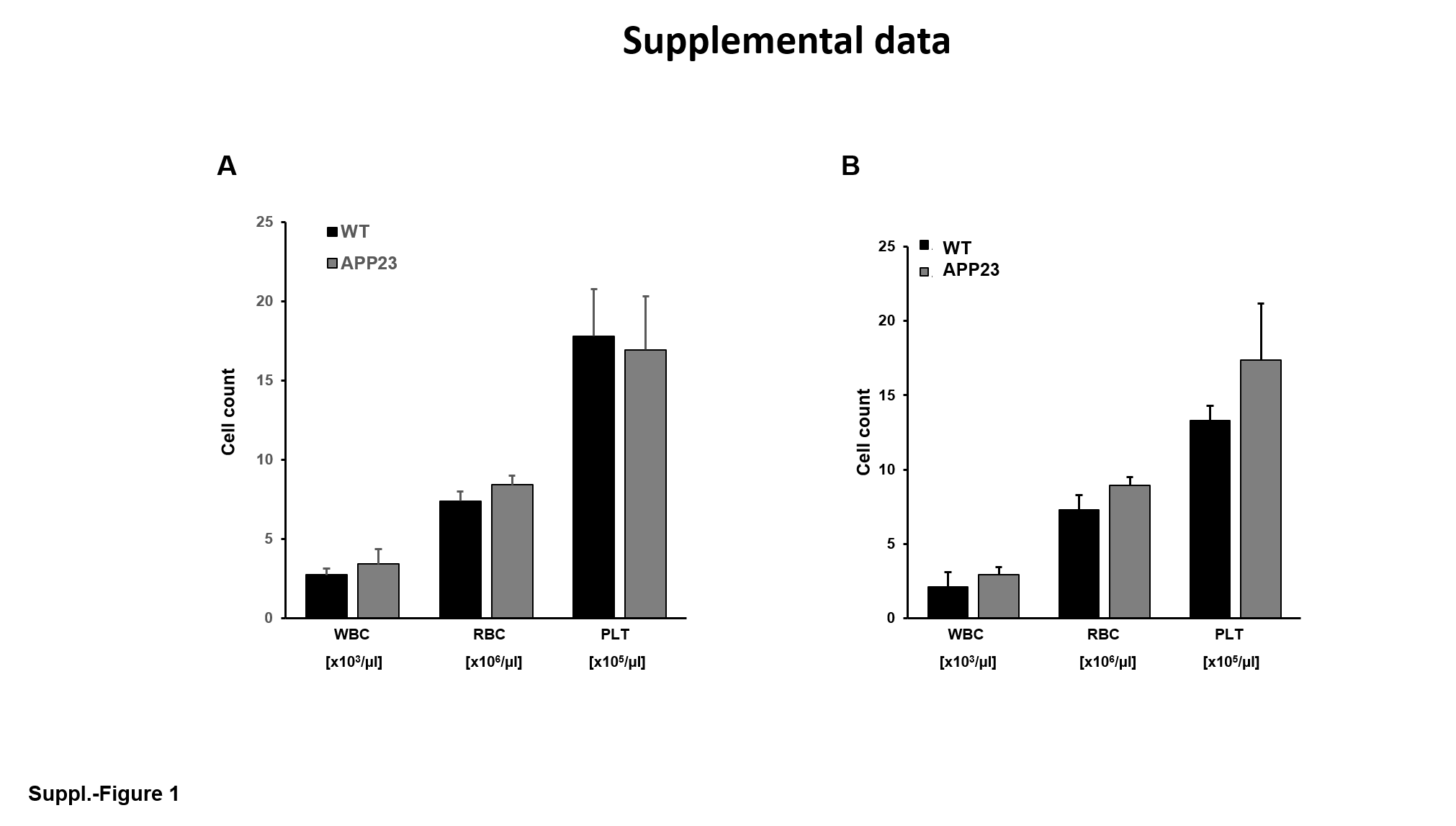
